# GEM-pRF: GPU-Empowered Mapping of Population Receptive Fields for Large-Scale fMRI Analysis

**DOI:** 10.1101/2025.05.16.654560

**Authors:** Siddharth Mittal, Michael Woletz, David Linhardt, Christian Windischberger

## Abstract

Population receptive field (pRF) mapping is a fundamental technique for understanding retinotopic organization of the human visual system. Since its introduction in 2008, however, its scalability has been severely hindered by the computational bottleneck of iterative parameter refinement. Current state-of-the-art implementations either sacrifice precision for speed or rely on slow iterative parameter updates, limiting their applicability to large-scale datasets. Here, we present a novel mathematical reformulation of the General Linear Model (GLM), wrapped in a GPU-Empowered Mapping of population Receptive Fields (GEM-pRF) software implementation. By orthogonalizing the design matrix, our approach enables the direct and fast computation of the objective function’s derivatives, which are used to eliminate the iterative refinement process. This approach dramatically accelerates pRF estimation while maintaining full accuracy. Validation using empirical and simulated data confirms GEM-pRF’s accuracy, and benchmarking against established tools demonstrates an order-of-magnitude reduction in computation time. With its modular and extensible design, GEM-pRF provides a critical advancement for large-scale fMRI retinotopic mapping. Furthermore, our reformulated GLM approach in combination with GPU-based implementation offer a broadly applicable solution that may extend beyond visual neuroscience, accelerating computational modelling across various domains in neuroimaging and beyond.

## 1 Introduction

Within neuroscience, the research domain of retinotopic mapping is dedicated to unravelling the relationship between the visual field and the visual cortex. Since early observations in the 1900s suggested a direct mapping between the visual field and visual cortex (Holmes, 1918; Tatsuji, 1909), many researchers have engaged in the development of techniques for assessing the retinotopic features of the visual system (for a review see (Wandell & Winawer, 2011)). Notably, functional magnetic resonance imaging (fMRI)-based techniques have become pivotal for non-invasive retinotopic experiments based on travelling waves stimuli (DeYoe et al., 1994; Engel et al., 1994). Numerous studies have established a functional topographic mapping of the human visual system using Positron-Emission Tomography (PET) (Fox et al., 1987; Zeki et al., 1991) and fMRI (Schneider et al., 1993). In particular, (Sereno et al., 1995) proposed mapping the retinotopic organization of multiple visual areas on the cortical surface based on phase information in fMRI data using periodic visual stimulation. Taking into consideration the eccentricities and polar angles of the corresponding receptive fields, they were able to functionally segment visual area regions such as V1, V2, VP, V3, and V4.

At the beginning of the twenty-first century, a new mathematical model for retinotopic analysis was proposed, referred to as population receptive field (pRF) mapping (Dumoulin & Wandell, 2008). Unlike previous methods, this approach is no longer limited to periodic stimulus designs and thus allows the use of arbitrary visual stimuli.

Numerous studies have used the pRF approach for visual field mapping (Benson et al., 2022; Bridge et al., 2023; Elul & Levin, 2024; Groen et al., 2022). In addition, applications in healthy participants with simulated visual field deficits (Haak et al., 2012; Hummer et al., 2018; Linhardt et al., 2022) and in clinical populations have demonstrated the potential of pRF mapping for staging retinal pathologies and tracking retinal disease progression (Molz et al., 2023; Prabhakaran et al., 2021; Ritter et al., 2019, 2024). These studies collectively enhance our understanding of visual field mapping in both simulated and actual vision loss. Another application of retinotopic mapping is the decoding of visual information, which is crucial for developing brain-computer interface (BCI) devices. These devices could one day reliably reconstruct cortical activity to the visual field, from reconstructing perceived or imagined letters (Senden et al., 2019) to complex visual scenes (Takagi & Nishimoto, 2023).

As pRF applications evolve, algorithms are needed that can provide swift retinotopic estimation results from even larger datasets. In the current implementations, pRF mapping techniques involve computing predicted time courses for each pRF position in the visual field. These time courses ultimately depend on the visual stimulus used in the experiment and the chosen pRF model, such as the isotropic 2D Gaussian (Dumoulin & Wandell, 2008) or the difference of Gaussians (Zuiderbaan et al., 2012). The algorithm then identifies the best-fit model signal for the measured fMRI time course in every voxel to estimate the pRF parameters spatial position (µ_x_, µ_y_) and other chosen model’s parameters (such as size (σ) in case of 2D isotropic Gaussian model). This fitting procedure uses optimization algorithms to minimize the difference between the predicted and measured fMRI time courses. It typically involves a two-stage process where initial coarse fitting is followed by refined fitting. Notably, the refined fitting procedure is commonly built as an iterative, computationally demanding optimization process.

Several implementations for such pRF mapping-based approaches are currently used by researchers such as mrVista (Dumoulin & Wandell, 2008) and SamSrf (D. Sam Schwarzkopf, 2018). These tools perform all computations on the CPU. There are other pRF mapping implementations such as DeepRF (Thielen et al., 2019), fast-pRF (Bhat et al., 2021) and qPRF (Waz et al., 2025), which specifically aim to minimise computational times. The DeepRF method is a deep learning-based approach and requires model training to predict the pRF parameters estimation for the measured fMRI data. Training the deep learning model for the DeepRF approach takes several hours and each specific experiment setup requires a model trained with the same setup. Given the long training time and the necessity for identical experimental setups between test and training data, this pRF mapping implementation might not be optimal for scenarios requiring frequent configuration changes. On the other hand, the fast-pRF method makes the pRF mapping procedure extremely fast on the CPU, however, it suffers from a loss in fitting precision. Therefore, it is only suitable for applications where accuracy can be sacrificed in favour of faster execution times, as e.g. real-time applications. Lastly, the qPRF method uses a searchable similarity-based tree to refine pRF parameters efficiently. By storing neighbor relationships, it enables rapid refinement. However, tree construction takes several hours, which may be a consideration for studies requiring frequent reconfiguration, such as comparative analyses with varying stimuli.

Considering the currently available toolboxes, we have identified a gap in pRF mapping implementations for applications that require highly accurate estimations combined with minimal computation times. To address this gap, we developed a novel pRF analysis software from scratch that (1) takes full advantage of the computational capabilities of current Graphics Processing Units (GPUs) and (2) optimises the mathematical procedure for time course fitting by reformulating the algorithm.

The utilisation of GPUs began in the late 1990s but was mostly limited to graphics applications. Their usage for high-performance computing in the framework of General-Purpose GPU (GPGPU) programming emerged with the introduction of the Compute Unified Device Architecture (CUDA) libraries, released in the mid-2000s. This marked a substantial milestone in facilitating GPGPU programming and sparked widespread adoption of the use of GPGPU hardware. Since then, numerous research fields have benefited from the unique computational capabilities of GPU accelerated software (Dally et al., 2021; Mišić et al., 2012), particularly in machine learning (Thielen et al., 2019).

For pRF analysis, however, the most commonly used software packages do not take full advantage of the computational power provided by GPUs. This is partly because the mathematics involved in pRF mapping are not easily transferred to the parallel processing paradigm of GPUs. Typically, the fMRI data processing steps utilise a general multivariate regression model, also known as General Linear Model (GLM) (Poline & Brett, 2012). In the case of pRF mapping, as proposed by (Dumoulin & Wandell, 2008), the application of GLM is required in both coarse as well as refine-fitting steps. However, during the refine-fitting process, the GLM needs to be applied iteratively to find the pRF parameter estimations. These requirements have made it challenging to implement a GPU-based pRF analysis application as GPUs are best suited for applications involving Single Instruction, Multiple Data (SIMD) processing.

In this study, we propose a novel implementation of pRF analysis based on a modified linear regression method, which can organise the data operations to follow a SIMD pattern. Our approach of GPU-Empowered Mapping of pRF (GEM-pRF) is encapsulated within a Python-based software package using CUDA libraries. To optimise efficiency and enable easy exploration of various setups, our methodology specifically employs the CuPy package-an open-source Python wrapper for CUDA (Okuta et al., 2017). Specifically designed for GPU operations, it follows NumPy’s API structure and includes the CUDA C++ runtime library NVRTC (Raschka et al., 2020), providing a Python API for compiling and executing CUDA kernels at runtime. This key feature enables the integration of native CUDA kernels (implemented in C/C++) to accommodate diverse pRF modelling approaches. We demonstrate the accuracy and applicability of GEM-pRF on both simulated and empirical data to show how GEM-pRF enables high-accuracy pRF analyses with greatly reduced processing time.

## 2 Methods

### 2.1 pRF modelling and fitting

The general idea behind pRF mapping is to compute expected time courses for the BOLD (blood-oxygen-level-dependent) signal based on a pRF model, a stimulus aperture, and a hemodynamic response function (HRF). The stimulus data depends on the experimental paradigm and must be provided as input for the analysis. As the HRF reflects a neurophysiological phenomenon, it varies between subjects. While some software implementations allow the computation of a subject-specific HRF, our software is designed to use either user-defined HRF values or a standard HRF as input. Consequently, while the stimulus and the HRF are typically assumed constant across voxels for a given experiment and subject, the pRF model depends on several parameters that must be estimated on a voxel-by-voxel basis. Following a similar methodology to the pRF fitting procedure outlined by (Dumoulin & Wandell, 2008), our implementation adopts a coarse-to-fine fitting strategy to estimate the pRF parameters for each measured fMRI time course. However, our approach introduces a novel streamlined linear regression method, optimised for high performance through GPU acceleration.

In classical pRF modeling, a pRF is represented as a function of spatial position in the visual field, denoted by *μ_x_*and *μ_y_*, which corresponds to the center coordinates of the receptive field. Additional parameters, such as the receptive field size σ, may also be included depending on the chosen pRF model. For instance, a commonly used model is based on a 2D Gaussian function, where σ defines the isotropic standard deviation of the Gaussian receptive field.

Mathematically, a pRF can be expressed as a function *f_ij_*(**θ**) that depends on a set of *K* parameters, where **θ** ∈ ℝ*^K^*, while the indices *i* and *j* represent the spatial positions in the visual field. In this paper, we focus on the 2D Gaussian pRF model with *K* = 3 parameters, which leads to **θ** = (*μ_x_, μ_y_*, σ). However, more complex models incorporating additional parameters can also be implemented in the GEM-pRF framework.

Another essential aspect in computing predicted time courses is the stimulus aperture, which determines the neuronal response patterns elicited during a pRF mapping experiment. The stimulus aperture can be mathematically defined as a three-dimensional matrix:

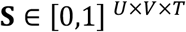

where:

- *U* × *V* represents the spatial dimensions of the stimulus grid,
- *T* is the number of time points in the experiment,
- Each element of **S** takes a value between either 0 (no stimulus) or 1 (stimulus present).

When a visual stimulus appears at a particular location and time, it triggers neural responses. These responses are typically modeled using the HRF, denoted as *h*(*t*), which describes the linear response of the BOLD signal to a neural event. Given this, a pRF model time course **p**(**θ**) corresponding to a particular parameter combination **θ** can be computed as follows:

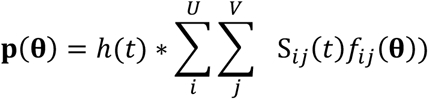

However, considering the linearity of convolution operation, we can first convolve the given stimulus **S** with *h*(*t*) and then use the HRF-convolved stimulus 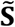 version to compute the model time courses. This can be represented as:

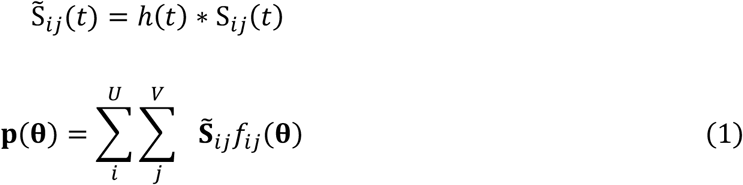

For pRF mapping analysis, the general linear model (Friston et al., 1995) is used. This model uses linear least squares optimization with a design matrix including effects of interest and additional confounds to model the task-based BOLD effect. The design matrix **X**, with a model time course **p**(**θ**) (i.e. P(**θ**, t) from equation (1)) for a particular modelling parameter combination **θ**, and the additional regressor matrix **R** can therefore be represented as:

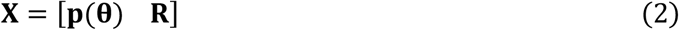

The regressor matrix **R** may contain different nuisance regressors and low-frequency functions, but, for the purpose of this work, must at least contain a constant term. Based on ordinary least square fitting, the residuals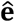between a measured time course **y** and a prediction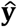can be represented as:

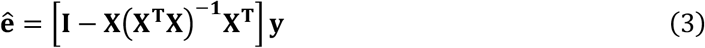

where **I** represents an identity matrix.

Since the prediction is invariant with respect to a change of basis in the column space of the design matrix, we can replace the design matrix with an orthonormal version. We are assuming the regressor matrix **R** to be already orthogonalized (e.g. by using Gram-Schmidt or QR decomposition) and then explicitly orthogonalizing each pRF model time course **p**(**θ**) with respect to **R**. The result is then normalised and denoted as **p**^′^(**θ**), which represents a prediction time course. This results in an orthonormalized regressor matrix **X^′^**, given as:

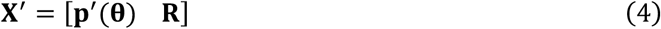

Upon substituting the modified regressor matrix **X’** in the equation (3) for residual errors, and solving it for the residual sum of squares (RSS), we obtain:

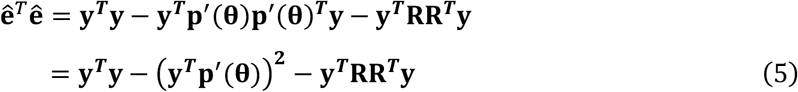

In pRF mapping, the ultimate goal is to compute the parameter **θ** that minimises the residual sum of squares (RSS). Upon careful examination of the new RSS equation (5), it becomes evident that both **y*^T^*y** and **y*^T^*RR*^T^*y** are independent of any modelling parameters **θ**. Therefore, the RSS is dependent on a prediction time course solely through the term (**y*^T^*p**^′^(**θ**) **^2^**.

Since (**y*^T^*p**^′^(**θ**))**^2^** directly affects the RSS, maximizing this value corresponds to finding the prediction time course that minimizes the residual error. This term represents the square of the dot product between **y** and **p**^′^(**θ**), and its value is maximized when both the time courses are either correlated or anti-correlated. In the case of anti-correlation, the dot product would yield negative values.

However, the goal is to find a prediction time course **p**^′^(**θ**) that correlates highly with the fMRI time course **y** (indicating a strong positive correlation). Thus, to focus on maximizing the correlation (and avoiding negative values due to anti-correlation), we can safely drop the square. With this observation, we can define the objective function 𝒞(**θ**) as the dot product between **y** and the prediction time course **p**^′^(**θ**), as shown below:

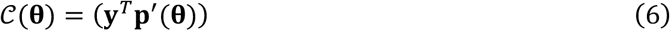

With the above-defined objective function, the coarse fitting step tries to maximize the objective function value.

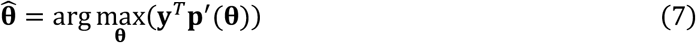

#### 2.1.1 Coarse fitting

The above reformulation of the objective function (equation (6)) especially simplifies the process of determining the best-fitting parameter combination from a given set of parameter combinations. This is crucial for determining an initial combination in the vicinity of the global optimum, here denoted as coarse fitting, which is then refined in a subsequent optimisation routine.

Given a set **Θ** of *N* parameter combinations **θ**, the best fitting parameter coarse parameter combination 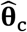 for a measured time course **y** in the coarse fitting step is therefore:

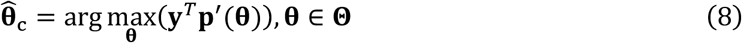

This can be efficiently implemented on a GPU, or in other parallelized systems, by constructing matrices from the prediction, as well as the measured time courses and performing a matrix multiplication, followed by an argmax operation. In GEM-pRF, this is implemented on the GPU using custom CUDA kernels in C/C++ alongside Python’s CuPy library.

It is worth noting that similar to the pRF mapping method proposed by (Dumoulin & Wandell, 2008), our method also uses a linear least square approach. However, our proposed method of using an orthogonalized design matrix eliminates the need for computing beta weights explicitly. This effectively reduces the computational load and enables the explicit computation of derivatives of the RSS objective function which is used for improving the initial coarse-fitting solution.

#### 2.1.2 Refine fitting

After the initial coarse fitting, where the optimal pRF parameters are selected from a discrete sampling space **Θ**, a fine-fitting step is performed to refine these estimates. Given a sufficiently dense sampling space **Θ** and a smooth objective function 𝒞(**θ**), the global maximum is expected to be in the vicinity of the coarse-fit solution. In our implementation, the refinement step efficiently determines this maximum without requiring iterative optimization.

##### Quadratic Approximation for Refinement

Traditional pRF mapping methods are based on iterative refinement of coarse estimates for each measured fMRI signal **y**, which is computationally expensive. Instead, GEM-pRF takes advantage of prediction time courses and their derivatives available on GPU side to compute the derivatives of the objective function directly, thus reducing overall computation times. This enables a non-iterative refinement based on a quadratic approximation of the residual sum of squares (RSS) values in the local neighbourhood of the coarse-fit parameters as illustrated in Figure 2.

Denoting the estimated coarse-fit parameters as 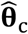, we consider the objective function values in their local neighborhood **Θ***_N_*⊆ **Θ**. Here, *N* represents the set of neighbours around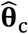 in the multidimensional discrete sampling space **Θ**. These objective function values can be approximated using a multidimensional quadratic function:

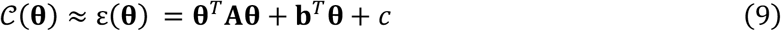

derivatives in the neighborhood
where:

- **A** is a symmetric square matrix
- **b** is a vector,
- *c* is a scalar value.

**A, b** and *c* coefficients in the above equation (9) are estimated using a linear least-squares fit to the function values 𝒞(**θ**) and their partial derivatives in the neighborhood of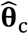 for each parameter combination **θ** using,

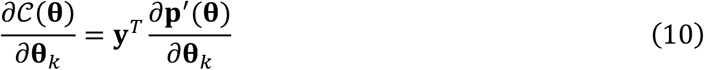

The first-order derivative of *ε* from equation (9) with respect to **θ** is given by:

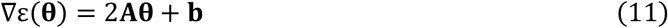

And by equating the above equation (11) to zero (i.e. ∇ε(**θ**) = 0), we obtain the following equation that can be solved to obtain refined-fitting parameters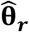:

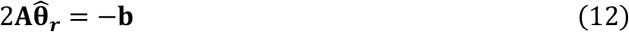

This refined solution 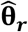 is accepted if its function value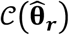 (from equation (6)) is greater than that at the coarse-fit estimate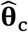.

Refinement relies only on a local neighbourhood, and the neighbours for each pRF parameter remain static. As a result, the neighbours and their corresponding design matrix for quadratic approximation are either precomputed or can be computed in parallel with the prediction signal computation. Furthermore, the neighbourhood data consists of only a few values, depending on the selected pRF modelling approach. This allows for quick processing on the CPU. Therefore, our overall refined fitting step efficiently takes place on the CPU side, as illustrated in Figure 1.

**Figure 1.**
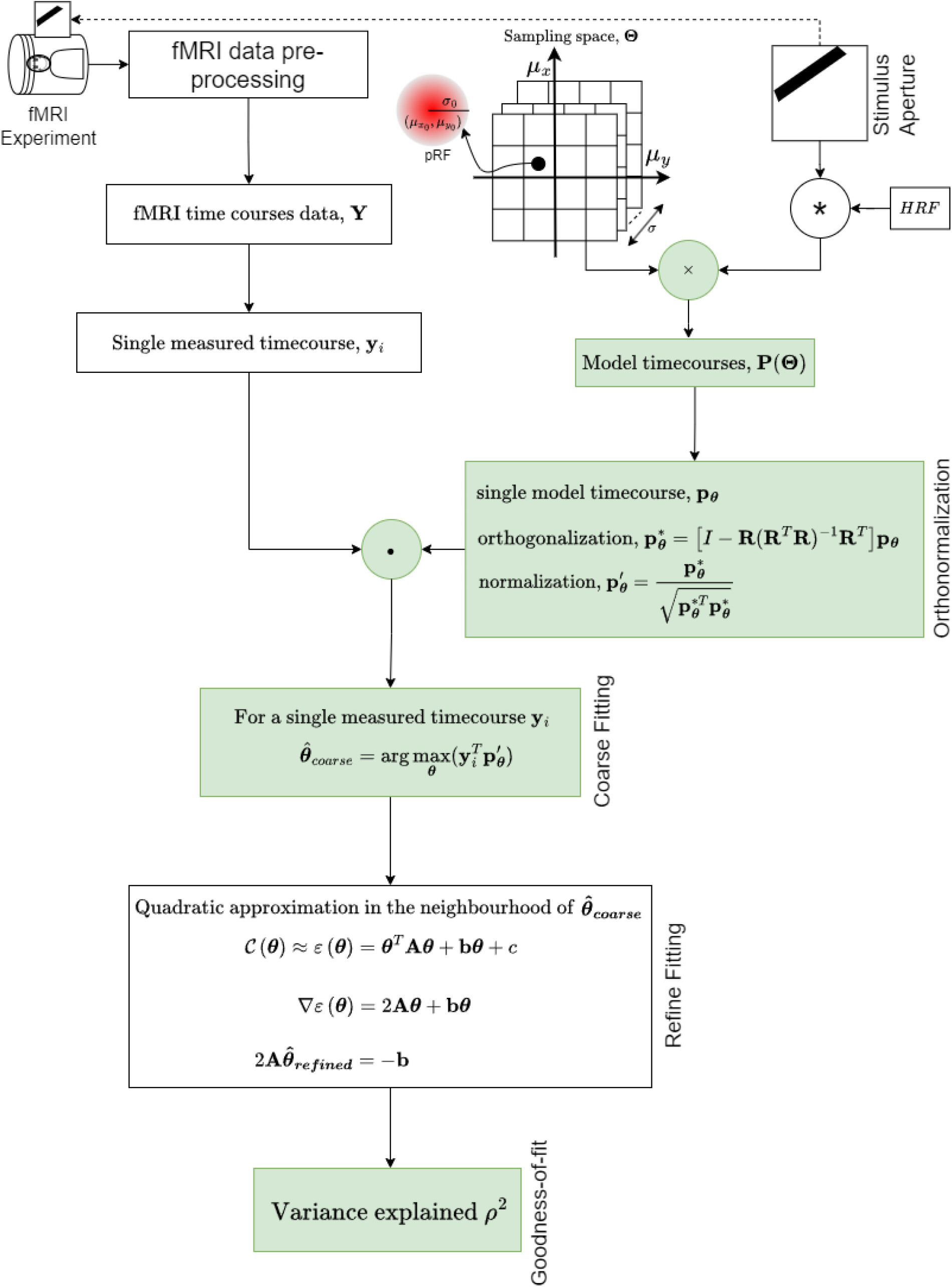
An illustrative flow chart representing the matrix dot product-based coarse-fitting step and a non-iterative refinement step. The green highlighted colours represent that the computation takes place on GPU for performance gain.

**Figure 2.**
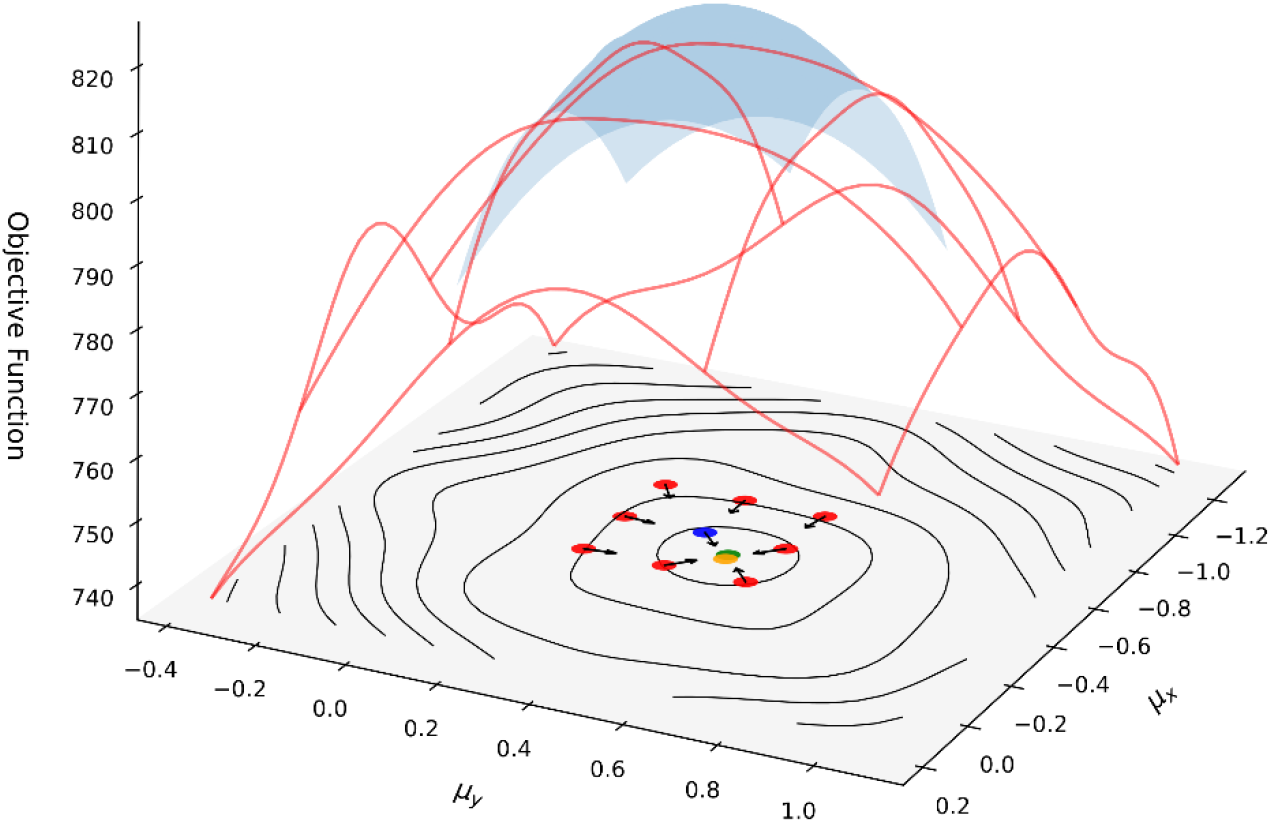
Illustration of quadratic surface approximation ε(**θ**) of the objective function 𝒞(**θ**) (equation (6)). The blue dot marks the spatial position of the coarse-fit result parameters (*μ_x_, μ_y_*, for a given σ). The red dots represent the immediate neighbours of the coarse-fit point in the sampling space. The red wireframe depicts the objective function values in an extended neighborhood around the coarse-fit parameters. The 3D translucent blue surface shows the refined quadratic approximation ε(**θ**) (equation (9)) near the coarse-fit (blue dot). The black arrows on the blue and red dots represent the gradients, where the arrow directions point towards the refined-fitting point. The green point represents the optimal parameters computed using non-linear iterative optimization function, while the yellow point represents the refined parameters obtained through our quadratic surface approximation. Notably, this example demonstrates an extreme case of a small pRF (small σ), leading to non-smooth objective function values in the extended neighborhood.

By employing this non-iterative, quadratic approximation approach, GEM-pRF significantly reduces computation time while maintaining accurate parameter estimations.

### 2.1.3 Variance explained

Similar to the original pRF mapping procedure proposed by (Dumoulin & Wandell, 2008), we incorporate a goodness-of-fit criterion to assess the quality of our predictions. This criterion quantifies how a prediction time series, generated using refined estimations of pRF parameters, aligns with a given measured fMRI time course, where all nuisance regressors were already removed. For this, we can compute the nuisance regressed time course **y**^*^ by using the equation,

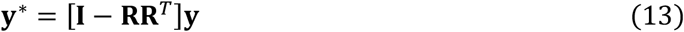

to compute the variance explained *ρ*^2^ as our measure of goodness-of-fit, calculated using the equation:

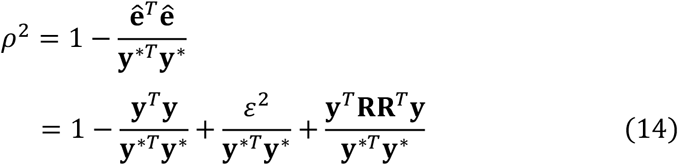

### 2.2 Multiple runs

The GEM-pRF implementation uses an approach equivalent to averaging for the joint analysis of multiple runs. To derive results for multiple runs, we continue to use our reformulated linear regression approach and extend it to the sum of the vector projections of the measured signals against the modelled signals for individual runs. Our approach can also jointly analyze the runs that are measured with different stimulus paradigms. For instance, if *M* numbers of distinct runs need to be jointly analysed, and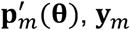 denote the orthonormal time courses and the measured fMRI time course for the *m*-th run respectively, then the pRF parameters for the joint analysis are obtained by identifying the parameter combination **θ** that maximises the sum of the products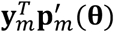 overall *m* with the objective function

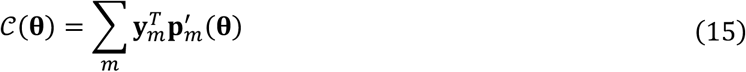

### 2.3 Sampling space

As mentioned above, the coarse-fitting step involves the definition of a parameter set **Θ**, which is used both for the coarse and the refined fitting process. Therefore, it is of utmost importance to choose a dense sampling for the pRF model’s parameters to create a set of representative parameter combinations. In the case of a 2D Gaussian pRF mode, the three parameters are the pRF spatial position *μ_x_, μ_y_*and size σ. While in its simplest form the combination of these spatial parameters can be formed by defining a uniform grid, some studies may employ more complex spatial distributions of the pRFs, such as hexagonal, circular, or other intricate shapes. The GEM-pRF program therefore provides the possibility to use complex distributions of the spatial parameters (*μ_x_*and *μ_y_*). Such custom spatial distributions can be provided as an input parameter. These combinations are then duplicated in other dimensions to generate the prediction time courses with all parameter combinations. Currently, for the isotropic 2D Gaussian model for pRF mapping, the default sampling space configuration of GEM-pRF is 151×151×16, corresponding to the parameters *μ_x_, μ_y_*and σ. Notably, the sampling space extends beyond the actually stimulated visual field. For instance, if the experimental stimulus aperture has a radius of 9°, the sampling space is defined for an extended visual field of 13.5°.

### 2.5 Data transfer considerations

The total number of model signals depends upon the defined pRF sampling space and commonly spans several thousand, entailing memory necessities ranging from several hundred megabytes to a few gigabytes (with 64-bit precision). To process the data on the GPU (referred to as device), the data must be present on the device. Such transfers of data from the CPU (referred to as the host) to the GPU device and vice-versa are predominantly facilitated by the Peripheral Component Interconnect-Express (PCIe) high-speed bus standard (Gorelick & Ozsvald, 2020), especially on the low-end consumer computing systems. The bandwidth provided by PCIe is critical for the performance of GPUs, especially in scenarios where large amounts of data need to be transferred between the CPU and GPU. Therefore, GEM-pRF tries to minimise such transfers, for example, the model signals are directly computed on the GPU side and kept there until the analysis finishes. Figure 1 provides an overview of various computation steps and the requirement of data availability on either the CPU or GPU side.

### 2.6 Multi-GPU environment

In a multi-GPU environment, GEM-pRF distributes the various computationally expensive and GPU memory-demanding execution steps onto different GPUs. This capability makes it possible to run the pRF analysis with a much larger number of model signals and therefore specifying denser pRF sampling space. For a multi-GPU cluster, the GPUs that are available for processing can be specified so that the program can automatically distribute memory requirements uniformly on all the specified GPUs. Therefore, all specified GPUs must have enough memory available for processing.

### 2.7 Data

To evaluate the results of pRF parameter estimation obtained using GEM-pRF, we employed both simulated and empirical data. For assessing computational time, we furthermore utilised a public dataset acquired at 7T and a locally acquired dataset on 3T scanner.

### 2.7.1 Simulated data

Simulated fMRI time series data was generated using the prfsynth docker image (version d9b64480cf1e) provided by a publicly available validation framework (Lerma-Usabiaga et al., 2020). To assess the accuracy of our implementation, we generated simulated fMRI time courses for three spatial locations (P, Q, and R) in the visual field. As shown in Figure 3(a), the parameters for the simulated pRF were set as follows: (*μ_x_*= 0, *μ_y_*= 0, σ = 1) for location P, (*μ_x_*= 3, *μ_y_*= 3, σ = 1) for location Q, and (*μ_x_*= 6, *μ_y_*= 6, σ = 1) for location R. The stimulus is considered to be extending from -10 to +10 degree visual angle. The pRF positions are chosen at different eccentricities to assess the accuracy of pRF parameter estimations between the fixation centre and the periphery close to the stimulation border. The simulated data was generated at two different white Gaussian noise levels to evaluate the accuracy and robustness of our implementation. These noise levels were specified in the configuration file provided by the validation framework for the prfsynth docker image. In the low-noise condition, the data achieved an average variance explained value of 88%, while in the high-noise condition, the average variance explained was 47%. Using the validation framework (Lerma-Usabiaga et al., 2020), we generated 5000 simulated time courses for each position. The average estimated position and pRF size were calculated by averaging the estimation results from all 5,000 simulations. However, for clearer visualization of the distribution of the results, the panels (c) and (d) in Figure 3 only displays 50 randomly selected individual estimations for each case, represented by grey circles.

**Figure 3.**
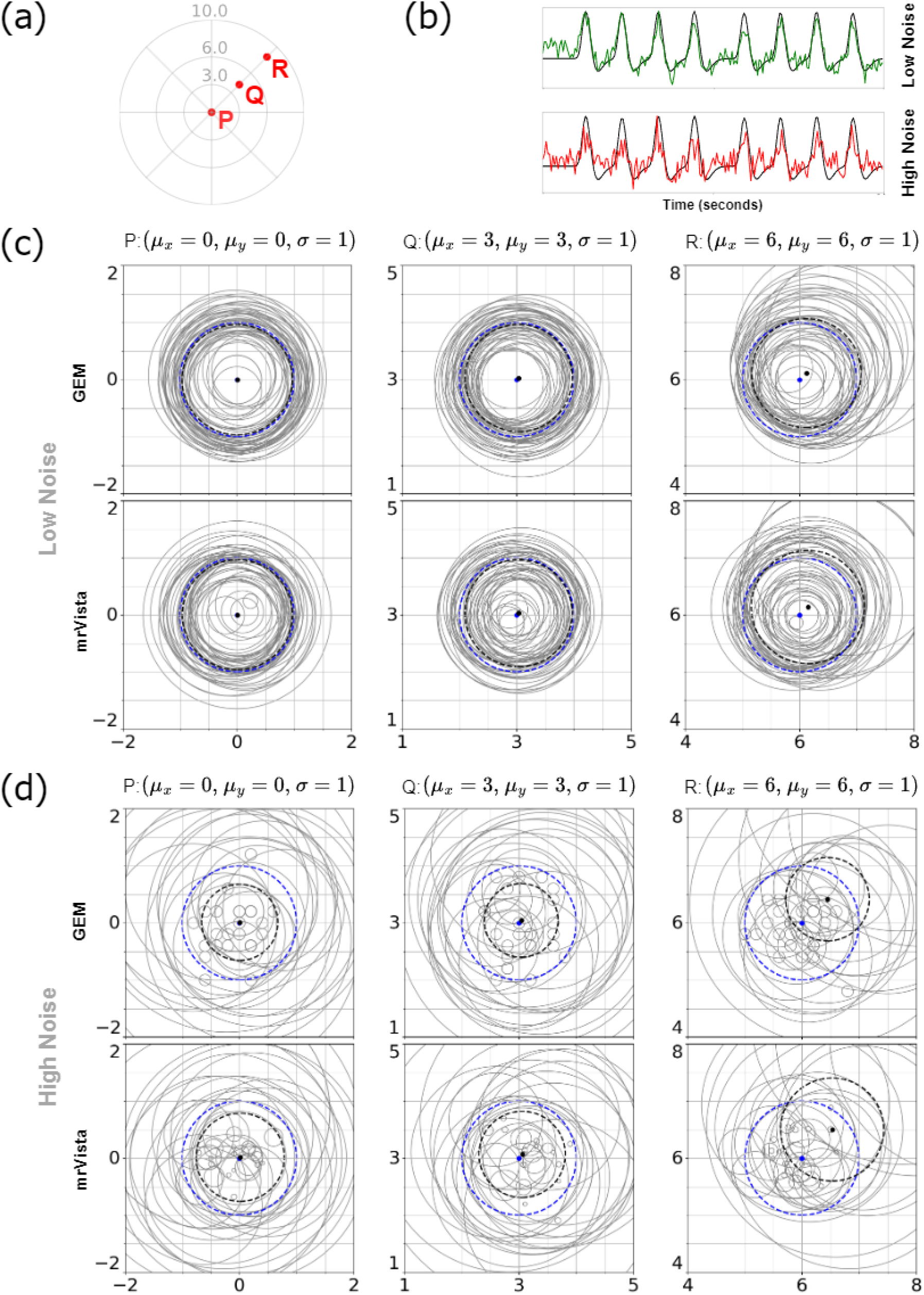
Comparison of population receptive field (pRF) estimation results at different eccentricities for ground truth data under low and high noise conditions. (a): Illustration depicting the spatial positions (P, Q, and R) of selected receptive fields used to generate simulated data. (b): Visualization of noise levels for low and high noise scenarios. (c, d): Comparison of estimation results using GEM-pRF and mrVista for low noise (Panel c) and high noise (Panel d) scenarios, generated for the selected spatial positions using pRF parameters P: (*μ_x_* = 0, *μ_y_* = 0, σ = 1), Q: (*μ_x_* = 3, *μ_y_* = 3, σ = 1), and R: (*μ_x_* = 6, *μ_y_* = 6, σ = 1). In Panels c and d, the blue dot and blue dashed circle indicate the ground truth pRF center and its size, respectively, while the black dot and black dashed circles represent the averaged estimated pRF center and its size.

To compute the pRF parameter estimation using GEM-pRF, we sampled the visual field across 151×151 spatial positions, spanning from -15.0 to +15.0 degrees, and duplicated these positions for 16 different pRF sizes linearly ranging from 0.1 to 5. This resulted in a total of 151×151×16 predicted time courses. And, to compare our estimation results with those of mrVista, we computed the mrVista pRF parameter estimations using the prfanalyze-vista Docker container (version 2.3.1_3.1.2) by (Lerma-Usabiaga et al., 2020), with its default configuration. This container is publicly available at GitHub (github.com/vistalab/prfmodel).

#### 2.7.2 Empirical data

For evaluation with empirical data, a healthy participant (25 yrs, female) was measured in a single session and 5 pRF runs were acquired using a 64-channel head coil on a 3T PrismaFit scanner (Siemens Healthineers, Erlangen, Germany). The study framework was approved by the local ethics board and adhered to the Helsinki Declaration. The full coil was used for anatomical measurements, while for the functional measurement, only the head coil’s lower part was used. Functional data was acquired using the CMRR EPI sequence (Moeller et al., 2010) with a spatial resolution of 1.5 mm isotropic and the following parameters: TE = 38 ms, TR = 1000 ms, multiband factor = 3, partial Fourier = 6/7, matrix size = 80x80, field of view (FoV) = 120x120 mm, phase encoding direction = left to right, flip angle = 55°, and slice spacing = 10%. Each run lasted 5 minutes, yielding 300 volumes, each consisting of 32 slices. The slices were aligned parallel to the calcarine sulcus, covering the participant’s occipital cortex. Anatomical imaging was performed using the MP2RAGE sequence (Marques et al., 2010) with a 1 mm isotropic resolution (TE = 2.98 ms; TR = 4000 ms, TI = 700/2500 ms, FoV = 256 x 216 mm, 160 slices and flip angle 4/5°).

For the fMRI runs, we employed a moving bar stimulus displaying a reversing checkerboard pattern, which was organized as a rectangular grid. The bar had a width of 1.2° visual angle, with a step size of 0.6° visual angle. The bar began moving from left to right, taking 36 steps for one crossing. The bar and the underlying checkerboard pattern rotated clockwise by 45° after each crossing. A single fMRI run consisted of eight different screen-crossing directions.

Following each diagonal crossing, a grey background image was displayed for 12 seconds as a baseline. The stimulus aperture had a diameter of 18°.

Data were preprocessed using fMRIPrep (version 23.1.4), which included slice-time correction, motion correction, and distortion correction. The pRF analyses were performed on data averaged over all five runs and masked for visual areas using prfprepare (Linhardt et al., 2025). For the empirical data analysis, the visual field was sampled in 151×151 spatial positions, ranging from -13.5° to +13.5° in the visual field, and duplicating these positions for 16 pRF sizes ranging from 0.5 to 5. The pRF estimation results were utilised to generate retinotopy maps (eccentricity, polar angle, and pRF size maps) for visual inspection using a custom in-house mapping software.

### 2.8 Performance analysis

To evaluate the computational times of the GEM-pRF implementation, we conducted performance tests on two platforms: a consumer laptop (ASUS ROG x13, AMD Ryzen 9, 8 cores, 3.30 GHz, 32 GB RAM, RTX 3050 Ti, 4 GB GPU Memory) and a High-Performance Computing (HPC) system (Intel Xeon E5-2698 v4, 40 cores, 2.20 GHz, 4x NVIDIA Tesla V100 DGX, 32 GB per GPU Memory). This setup provides a significant contrast for evaluating computational times on low-end and high-end systems.

To provide a comparison of computational times with respect to the size of an fMRI dataset, we created simulated fMRI datasets of different sizes (ranging up to 100,000 voxels) using the validation framework provided by (Lerma-Usabiaga et al., 2020). The ground truth parameters for all synthesized time courses were set to *μ_x_*= 0, *μ_y_*= 0 and σ = 1. Additionally, the white noise level in the validation framework was adjusted to achieve an overall variance explained of approximately 70%. To analyse the effect of the chosen configuration for the sampling space, we computed the pRF parameters estimation results with the following setups:

- GEM-pRF with sampling space 151×151×16 with refine-fitting enabled (our default configuration) on the HPC system
- GEM-pRF with sampling space 151×151×16 without refine-fitting on HPC system
- GEM-pRF with sampling space 151×151×16 without refine-fitting on consumer laptop
- mrVista, using its default configuration of the prfanalyze-vista container (version-2.3.1_3.1.2)

The above analyses allowed us to evaluate the computational performance across various dataset sizes and on systems with strong differences in their computing capabilities.

## 3 Results

All analyses were computed on either a standard laptop with a GPU or a high-performance computing system with multiple GPUs clusters (please refer to the Method section for details).

### 3.1 Simulated data analysis

The analysis of pRF parameter estimation by GEM-pRF accuracy relies on simulated fMRI data with added noise. This simulated dataset comprises 5000 fMRI time series for three distinct spatial positions. Since we possess the ground truth pRF parameters in this scenario, assessing the accuracy of our estimation becomes more straightforward.

In Figure 3(a), we illustrate the selected spatial positions (P, Q, and R) for which both low-noise and high-noise (as examples depicted in Figure 3(b)) simulated fMRI data were generated. Figure 3(c) & (d) present a comparison between the pRF parameters estimated by GEM-pRF, which are depicted by black dots, representing the average of individual estimations (grey circles, 50 randomly chosen samples). In the same plots, the ground truth values are represented by blue dots for spatial position and dashed blue circles for pRF size. The Figure also provides a comparison of the estimated parameters by GEM-pRF and the widely used tool mrVista. Analysing Figure 3(c), it becomes apparent that under the low noise scenario, the pRF parameters estimated by GEM-pRF and mrVista align closely with the ground truth for spatial positions P and Q. For the low noise scenario, minor deviations from the ground truth parameters can be observed for the peripheral position R. As further depicted in Figure 3(d), in the high noise scenario, on average both GEM-pRF and mrVista seem to reliably estimate the pRF parameters for positions P and Q. However, in this case, for both methods, notable deviations from the ground truth parameters can be observed for the peripheral spatial position R.

A closer examination of Figure 3(d), showing the estimation results, reveals several small grey circles by both GEM-pRF and mrVista. Since all the simulated fMRI time courses were generated with a ground truth pRF size of 1 (indicated by the black dashed circle), these small circles therefore suggest that the pRF sizes were underestimated by both implementations in the high noise scenario. This can also be observed in the high noise case shown in Figure 4.

**Figure 4.**
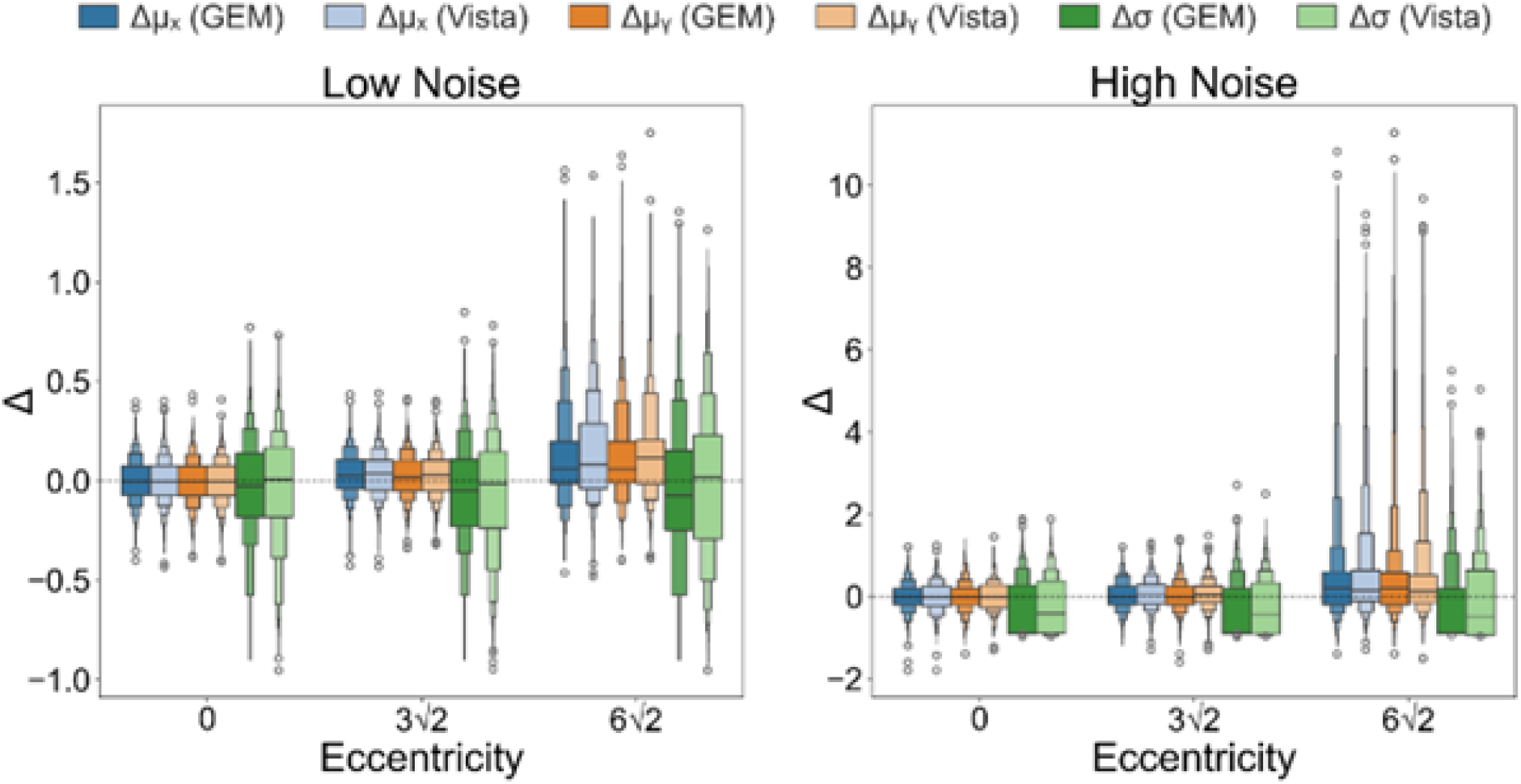
Comparison of GEM-pRF and mrVista for pRF parameter estimations using the simulated data across low-and high-noise scenarios. The Figure shows the differences between the estimated value and the ground truth for a given pRF parameter (i.e. *μ_x_, μ_y_* or σ). As indicated in Figure 3, the comparisons are conducted at the same spatial positions (P, Q, R) corresponding to eccentricity values of 0, 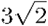 and 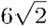.

The Figure 3(d) further reveals that the GEM-pRF results show several fixed size small circles and mrVista show even smaller circles.

Figure 4 provides a further analysis to compare the estimation results from the GEM-pRF and mrVista implementations. The boxenplots (Heike et al., 2017) illustrate the deviations (Δ) of the estimated parameter values from the ground truth. In the low-noise condition, both implementations showed minimal bias for position estimates (*μ_x_, μ_y_*), with small deviations centered around zero. However, mrVista exhibited greater variability in the size parameter (σ), particularly at higher eccentricities. In the high-noise condition, estimation errors increased across both software implementations. Additionally, at higher eccentricities, the variance of positional errors (Δ*μ_x_*, Δ*μ_y_*) was asymmetrically larger on the positive Δ side, suggesting a systematic bias towards the periphery. This pattern was more pronounced in the high-noise condition.

### 3.2 In vivo data analysis

We computed retinotopy maps using the default configuration of GEM-pRF, with 151×151×16 prediction signals for coarse fitting. The cortex overlays of the eccentricity, polar angle and pRF size maps are presented in Figure 5. Additionally, the coverage map, also shown in Figure 5, depicts a higher density of mapped pRFs near the foveal region compared to the parafoveal region. It further highlights a disparity in the number of mapped pRFs between the upper and lower vertical meridians.

**Figure 5.**
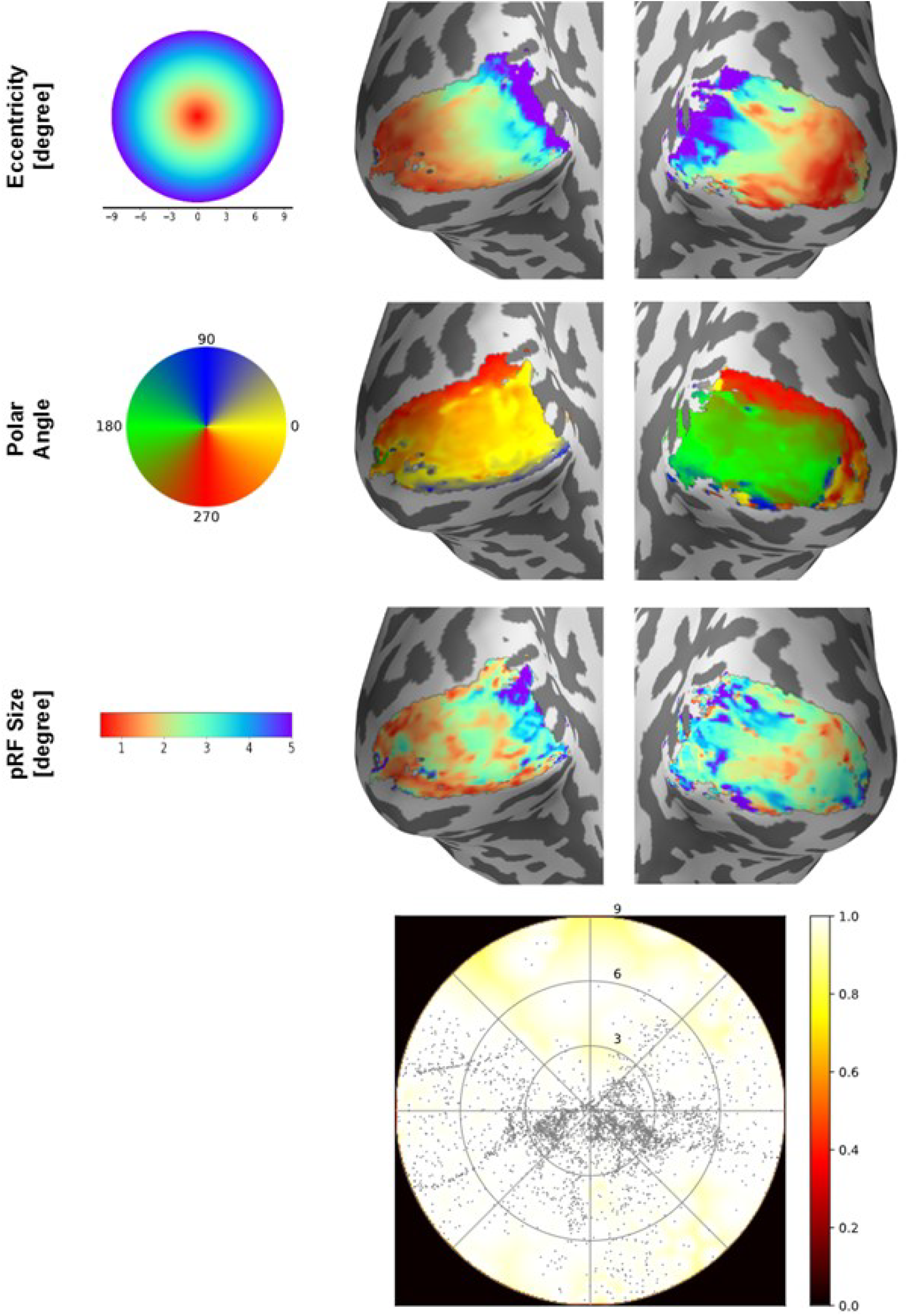
Visualization of estimated population receptive field (pRF) parameters overlaid onto an inflated cortical surface, delineating the left and right hemispheres of the brain. The bottom row displays an aggregate coverage map spanning both hemispheres. The pRF parameters were estimated utilizing the default settings of GEM-pRF, employing a grid of 151×151 pRF positions across the visual field (total diameter 18°), each replicated for 8 sigma values.

### 3.3 Performance analysis

Figure 6 presents a comparative analysis of the overall execution durations of GEM-pRF using two different pRF parameter estimation configurations: (1) using only coarse fitting and (2) using both coarse and refine-fitting. The total execution times include various stages of the analysis process, such as coarse fitting, refine-fitting (if enabled), calculation of variance explained, and writing results to a file. Additionally, depending on the specified sampling space and stimulus used, the software incurs some initialization time at the beginning, which involves the computation of prediction signals, the determination of neighbors for each pRF parameter set, and the generation of the corresponding design matrix for quadratic approximation.

**Figure 6.**
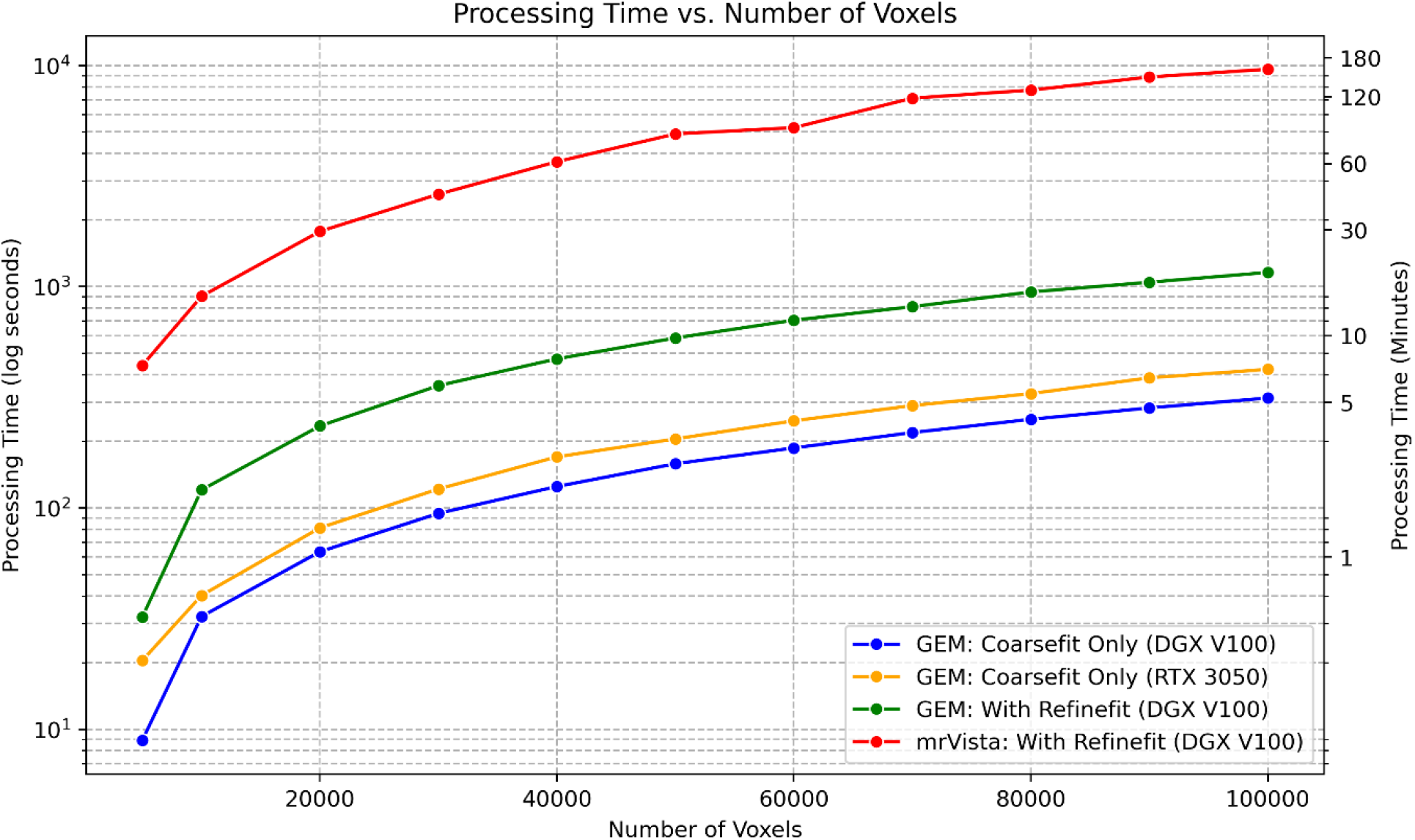
Comparison of mrVista vs. GEM-pRF computational times as well as the comparison of two GEM fitting configurations i.e. coarse fitting only and coarse fitting plus refine-fitting. The default parameter-sampling configuration is used i.e. a regular grid of 151×151×16. The computational times are depicted on a log scale.

In the current comparison scenario with the default sampling space of 151×151×16 and a bar visual stimulus of duration 300 seconds and size 101×101, the total initialization time was approximately 320 seconds on our HPC system. This initialization time is not included in the computational times shown in Figure 6 for two reasons. First, it is independent of the size of the fMRI dataset being analyzed. Second, the calculations performed during this initialization phase are required only once when analyzing multiple datasets. For instance, Figure 6 depicts the computational times required for datasets containing varying numbers of vertices (ranging from 5,000 to 100,000). In this case, the initialization computations were performed once and subsequently reused across all datasets. Furthermore, since the calculations that are part of the initialization times are static, they can be cached and can be loaded from cache to further reduce the initialization times.

A closer examination of Figure 6 reveals an almost linear increase in computational time with the size of the fMRI dataset. To evaluate the performance improvements achieved with GEM-pRF, we compared its computational efficiency against the widely used tool, mrVista. We conducted pRF analyses on simulated data using mrVista on our HPC system. As shown in Figure 6, mrVista’s default implementation required slightly less than 3 hours to perform pRF estimations on a dataset containing 100,000 voxels. In contrast, GEM-pRF demonstrated significantly faster processing times, completing the same task in approximately 6 minutes on the HPC system and around 7 minutes on a consumer laptop using only the coarse-fitting configuration (151×151×16 sampling space) and about 20 minutes on the HPC system when both coarse and refine-fitting were applied with the same sampling space.

## 4 Discussion

The proposed GEM-pRF implementation presents a novel approach to pRF mapping by utilising GPU acceleration for efficient processing of large fMRI datasets. Our implementation reformulates the GLM approach into projections on prediction time courses **p**^′^(**Θ**) which can be computed and evaluated on the GPU, enabling accelerated execution of the coarse-fitting stage of pRF estimation. We achieve this acceleration through a modified linear regression approach, facilitating SIMD data processing. Additionally, we introduce a novel method for refining coarse pRF estimated parameters as illustrated in Figure 2. For our refinement step, we assume that our objective function can be approximated by a quadratic function in the neighbourhood of the coarse fitting pRF estimation parameters. The maximum of the quadratic approximation is then used as the refined pRF parameters. These mathematical concepts are encapsulated within our GEM-pRF software implementation. Figure 1 outlines the core computation steps of the GEM-pRF software and illustrates the execution of these steps on the CPU and GPU. Furthermore, the GEM-pRF implementation provides an extensive configuration file, enabling researchers to specify input datasets in BIDS format and conduct pRF analysis with different settings for comparative analysis.

### Accuracy

The evaluation of pRF parameter accuracy utilising GEM-pRF was conducted through a dual approach involving empirical and simulated data. The cortex overlays of eccentricity, polar angle, and pRF size maps (Figure 5) exhibit expected retinotopic organization (M. M. Himmelberg et al., 2023; Wandell & Winawer, 2011) in an in vivo dataset. Eccentricity values increase from red to blue as we move from the posterior to the anterior end of the primary visual cortex (V1), aligning with known retinotopic organization. The polar angle maps follow the expected visual field representation, with the left hemisphere corresponding to the right visual field and vice versa. Furthermore, the coverage map shown in Figure 5 reveals more coverage along the lower vertical meridian as compared to the upper vertical meridian, which is also consistent with the previous studies (M. Himmelberg et al., 2021; M. M. Himmelberg et al., 2023). This empirical analysis validates the accuracy of our implementation for pRF analysis with the default sampling space configuration 151×151×16 with refine-fitting.

Additionally, quantitative analysis using simulated data demonstrated GEM-pRF’s ability to estimate pRF parameters for noisy fMRI data. However, estimations for peripherally situated receptive fields (e.g. Position R in Figure 3) showed larger deviations from ground truth, particularly under high-noise conditions, aligning prior research findings (Lerma-Usabiaga et al., 2020).

The high-noise case, as shown in Figure 3(d) and Figure 4, highlights an underestimation of pRF sizes by both implementations. At first glance, Figure 4 suggests that mrVista underestimates pRF sizes more than GEM-pRF. However, a closer look at Figure 3(d) reveals that while GEM-pRF estimates several small pRF sizes at a fixed lower bound, mrVista produces even smaller estimates. The small, fixed pRF sizes in GEM-pRF correspond to the lower bound of our specified sigma size for the given sampling space **Θ**, which was set to 0.1, as described in the methods section. Further analysis of these small pRF estimates showed that both implementations achieved similar average variance-explained (*ρ*^2^) values, which serves as a key goodness-of-fit metric for both methods, when applied to these simulated voxels. These findings suggest that while both GEM-pRF and mrVista exhibit large deviations in pRF size estimates for voxels with low-SNR time courses, they still produce equivalent *ρ*^2^ values, confirming their methodological consistency as expected.

Furthermore, the observed directional bias in position estimates at higher eccentricities suggests a systematic deviation in pRF localization, which may reflect an inherent limitation either in visual stimulation pattern or the fitting procedures of both methods. This effect warrants further investigation in future studies to determine whether it stems from methodological constraints or the choice of visual stimulation pattern. Understanding these biases will be crucial for improving pRF modelling, particularly in studies examining peripheral visual field representations.

### Speed

The GEM-pRF implementation provides a significant speedup, as depicted in Figure 6. The total computational time required for pRF parameter estimation analysis using our implementation on a large dataset comprising 100,000 voxels shows a speedup of one order of magnitude.

Figure 6 further demonstrates that the computational times on the consumer laptop system are higher than those on the HPC system for a sampling space of 151×151×16 without refined fitting. This discrepancy can be attributed to the fact that a significant portion of the computation involves matrix operations and objective function term evaluations, which are handled more efficiently by the DGX V100, likely due to its overall architectural optimizations for computational workloads. Furthermore, data transfer between the GPU and CPU contributes more to the overall runtime on the consumer laptop system, likely due to differences in PCIe bandwidth and memory access speeds. These factors collectively explain the increased processing time observed on the consumer laptop system equipped with the RTX 3050 GPU.

It is worth noting that the underlying mathematics for GEM-pRF implementation is a reformulated version of the originally proposed pRF mapping methodology by (Dumoulin & Wandell, 2008). Therefore, the pRF parameters and the retinotopic maps calculated with GEM-pRF are similar to those obtained by the mrVista gold-standard approach. This contrasts with some other previous implementations, which provide alternate methodologies for fast pRF mapping. For example, f-pRF (Bhat et al., 2021) provides the possibility for fast computation of pRF parameters estimation on a CPU. However, to favour speed over accuracy, they use tile coding and hashing based encoding of stimulus, without using typical GLM approach for parameter estimations. While this approach provides significant speed, it often leads to the mapping of parameters on the discrete grid values with significant deviations. Another technique that has also tried to address the speed issue for pRF mapping is the DeepRF implementation (Thielen et al., 2019). Deep learning-based methods hold great potential to reliably predict pRF parameters from measured fMRI data. However, their accuracy requires further independent validation. Additionally, the substantial training time needed for these models limits their suitability for research studies where it becomes necessary to train different models (e.g. for the comparison of pRF mapping results using different stimuli). There are also some other methods that have altogether tried to address the speed issue from a different perspective. For example, the deep learning-based implementation by (Ribeiro et al., 2021) returns estimated pRF parameters of the early visual areas without requiring functional scans. Their method predicts retinotopy maps using brain segmentation from anatomical scans using a geometric deep learning model trained on the Human Connectome Project (HCP) dataset (Benson et al., 2018). While this method significantly accelerates the mapping process, it comes with certain limitations. First, its accuracy is influenced by the reliability of the pRF estimations used for training, meaning biases introduced by biases introduced by stimulus selection in the experimental setup (Alvarez et al., 2015; Chang et al., 2025; Linhardt et al., 2021) could be learned by the deep learning model, potentially affecting the generalizability of its pRF estimations. Additionally, since the method relies solely on anatomical data to predict pRF parameters, it may not account for functional alterations due to visual pathway pathologies. Condensing these facts, our proposed GEM-pRF implementation provides the possibility for quick estimation of pRF parameters for large datasets (such as the HCP dataset), while maintaining high accuracy.

### Sampling space

As previously outlined, the refinement stage aims to enhance the accuracy of pRF parameter estimation achieved during the coarse fitting phase. This refinement process involves exploring parameter combinations within the vicinity of the initially computed coarse-fitting pRF parameters. These parameters, along with their neighbouring values, form a discrete set of combinations. The objective of refinement is to identify optimal parameter combinations lying between these discrete points, thereby better capturing the measured fMRI time course, **y**.

Given this rationale, selecting an adequately dense sampling space for pRF parameters is crucial for computing prediction time courses. Our default sampling space is set at 151×151×16, corresponding to the pRF parameters (*μ_x_, μ_y_*and σ). Additionally, the GEM-pRF implementation provides flexibility in utilizing various custom sampling configurations. It also accommodates complex, irregular spatial distributions of receptive fields. This adaptability enables the analysis and comparison of pRF estimation results across various spatial arrangements of receptive fields in the visual field.

### Joint analysis

It is customary in pRF analysis applications to perform a joint analysis of multiple datasets, obtained during different runs, sessions, or with different visual stimuli for a subject. The GEM-pRF package provides an out-of-the-box software implementation possibility to conduct such joint analyses.

### Scalability

In terms of scalability, our software program employs a batching procedure to handle the memory-intensive execution steps on GPUs. Since the estimation of pRF parameters for each voxel’s fMRI time series, **y**, is independent of others, the minimum batch size for processing measured fMRI data can even be reduced to a single voxel’s time series.

However, during the computation of coarse fitting, the vector projection of a voxel’s fMRI time series, **y**, is computed with respect to all the prediction time courses. Thus, in our current implementation, it’s required to hold all the prediction time series data in GPU memory. This necessitates a constraint wherein the user must select a sampling space **Θ** configuration such that the prediction time courses can be stored in the available GPU memory. Our tests have indicated that a sampling space of 151×151×16, corresponding to the pRF parameters (μ_x_, μ_y_and σ), achieves satisfactory accuracy and can be executed on consumer-grade GPU systems by disabling refine-fitting.

### Future scope of work

The chosen modular and generic development approach for the GEM-pRF implementation paves the way for future advancements. The software currently uses a 2D Gaussian model for pRF mapping, with the potential to incorporate alternative models, such as the Difference of Gaussian (Zuiderbaan et al., 2012), into the GEM-pRF open-source implementation. The GEM-pRF implementation already provides a framework of abstract classes, making it easy to integrate a new pRF mapping modelling approach.

Moreover, to enhance performance, further improvements can be explored, such as leveraging CUDA streams to achieve additional parallelization in CPU-GPU data transfer and processing. This optimization can lead to faster execution times and more efficient utilisation of computational resources, ultimately enhancing the scalability and usability of the GEM-pRF software.

## 5 Conclusion

We herein introduced a breakthrough solution that reworks the design matrix using orthogonalization, allowing us to compute the objective function and its derivatives directly on the GPU. This eliminates the need for iterative refinement, making the process significantly faster while keeping the accuracy unchanged. Our novel GEM-pRF analysis approach yielded accurate pRF mapping results in a fraction of the computation time required with the gold-standard mrVista software package. This increase in computational speed is due to a modified fitting procedure combined with GPU-powered acceleration and offers a modular and flexible approach for efficiently analysing even large fMRI datasets with varying configurations. Our evaluation demonstrated GEM-pRF’s accuracy in estimating pRF parameters, validated through both empirical and simulated data analyses. Notably, the software’s scalability allows for effective handling of memory-intensive computations, while its performance surpasses that of widely used tools, significantly reducing computation times without sacrificing accuracy. Looking ahead, GEM-pRF’s modular design provides possibilities for future enhancements, such as incorporating additional pRF mapping models and optimising CPU-GPU data processing. As the first GPU-accelerated implementation utilizing the traditional GLM-based fitting approach for visual field mapping, GEM-pRF offers a significant step forward in computational neuroimaging, enhancing the efficiency and accessibility of large-scale retinotopic mapping.

## CRediT authorship contribution statement

**Siddharth Mittal**: Data curation, Conceptualization, Methodology, Software, Validation, Visualization, Writing – original draft, Writing – review & editing. **Michael Woletz**: Conceptualization, Methodology, Validation, Visualization, Writing – review & editing. **David Linhardt**: Data curation, Conceptualization, Methodology, Validation, Visualization, Writing – review & editing. **Christian Windischberger**: Conceptualization, Methodology, Validation, Visualization, Writing – review & editing, Supervision, Project administration, Funding acquisition.

## Acknowledgements

This work was supported by the Austrian Science Fund (FWF; grant number: P35583).

## Declaration of generative AI and AI-assisted technologies in the writing process

During the preparation of this work the author(s) used Grammarly and ChatGPT in order to improve the readability of our manuscript. After using this tool/service, the author(s) reviewed and edited the content as needed and take(s) full responsibility for the content of the publication.

